# Pre-Clinical Blocking of PD-L1 molecule, which expression is down regulated by NF-κB, JAK1/JAK2 and BTK inhibitors, induces regression of activated B-cell lymphoma

**DOI:** 10.1101/502922

**Authors:** Christelle Vincent-Fabert, Lilian Roland, Ursula Zimber-Strobl, Jean Feuillard, Nathalie Faumont

## Abstract

Escape from immune control must be important in the natural course of B-cell lymphomas, especially for those with activation of NF-κB. The pre-clinical L.CD40 transgenic mouse model is characterized by B-cell specific CD40 signaling responsible for NF-κB continuous activation with a spleen monoclonal B-cell tumor after one year in 60% of cases. L.CD40 tumors B-cells expressed high levels of PD-L1. This expression was dependent on activation of either NF-κB, JAK1/JAK2 or BTK pathways since ex vivo treatment with the inhibitory molecules PHA-408, ruxolitinib and ibrutinib led to decrease of its expression. Treatment of L.CD40 lymphomatous mice with an anti-PD-L1 monoclonal antibody induced tumor regression with decreased spleen content, activation and proliferation rate of B-cells as well as a marked increase in T cell activation, as assessed by CD62L and CD44 expression. These results highlight the interest of therapies targeting the PD-1/PD-L1 axis in activated lymphomas with PD-L1 expression, with possible synergies with tyrosine kinase inhibitors.

## Introduction

Aberrant expression of the programmed death-ligand 1 (PD-L1, also known as B7-H1 or CD274) checkpoint molecule has been reported in many cancers such as breast, lung and colon tumors as well as during chronic viral infections like Epstein-Barr virus (EBV) infection for example (1,2). Efficacy of immunotherapies against the PD-1/PD-L1 axis in breast or colon cancers demonstrated the importance of the immune checkpoints in the control of emergence and growth of tumors (2). As reviewed recently, various publications have indicated that disruption of immune checkpoints is also a critical step in B-cell non-Hodgkin’s Lymphomas (NHL) (3). NF-κB, one of the most cited transcription factor in B-cell lymphomas, is able to increase tumor cell expression of PD-L1 either directly or indirectly (3). NF-κB constitutive activation is found either in aggressive diffuse large B-cell lymphomas (DLBCL) with an activated phenotype (ABC-DLBCL), or in indolent B-cell lymphomas such as chronic lymphocytic leukemia, Waldenström Macroglobulinemia, marginal zone B-cell lymphomas (MZL) (4). Here, we wanted to explore the putative interest of PDL-1 immune therapy against B-cell lymphoma with NF-κB activation. To experimentally address this question, we used a transgenic mouse model which specifically express in B-cells a chimeric protein composed of the transmembrane moiety of the Epstein Barr Virus latent membrane protein 1 (LMP1) and the transduction tail of CD40 (L.CD40 protein), that results in continuous activation of NF-κB, responsible for a spleen monoclonal B-cell tumor (L.CD40 B-cell lymphoma) after one year in 60% of cases (5).

## Results

Our previous transcriptome studies from L.CD40 mice suggested that those tumors might express high levels of CD274/PD-L1 (6). We thus analyzed the Immune Escape Gene Signature published by C Laurent *et al* (Supplemental Table 1) (7) from the Affymetrix transcriptome (HT MG-430 PM Array, Supplemental Tables 2 and 3) of a series of six L.CD40 B-cell lymphomas that were compared to their CD19_Cre littermate. As shown in Figure 1A, both PD-L1 (red arrow) and PD-L2 (orange arrow) were co-clusterized with the immunosuppressive interleukin 10 (IL-10, green arrow), CD80 and MCL1, being over expressed. Indeed, L.CD40 lymphoma cells from 12 months old mice expressed higher levels of PD-L1 than B-cells from age related CD19_Cre mice (Figure 1B).

**Figure 1.**
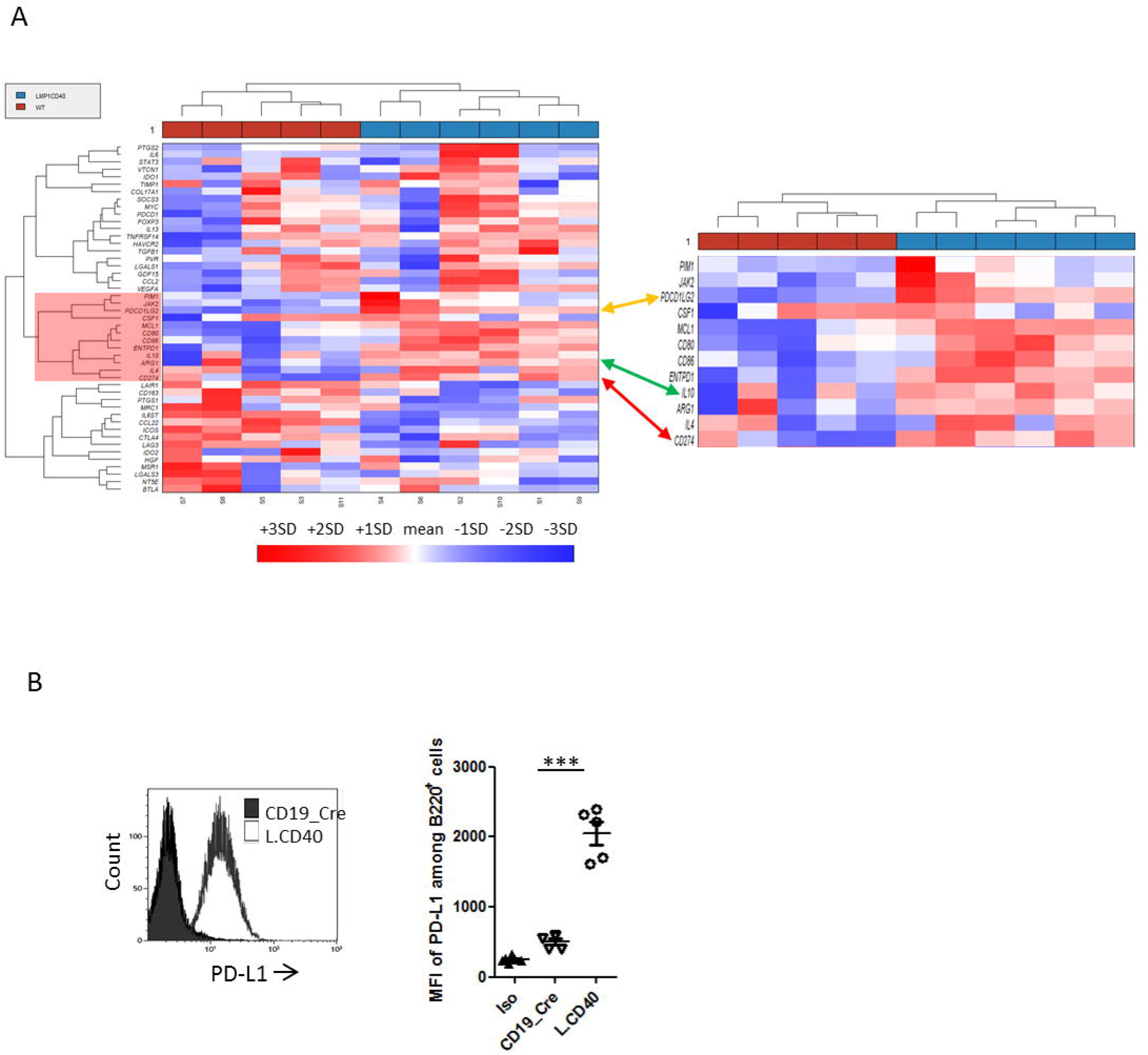
(A) Clustering of genes from the Immune Escape Gene Signature published by C Laurent *et al*. for L.CD40 and WT mice. The specific cluster for L.CD40 mice is highlighted in red. PD-L1 is pointed by the arrow. (B) Left panel, overlay example of PD-L1 monoparametric histograms gated on B220 B-cells. Right panel, flow cytometry Mean Fluorescence Intensity (MFI) of PD-L1 on B-cells from 12 months old control CD19_Cre and transgenic L.CD40 mice. Statistical significance was determined by unpaired t-test (***P<0.001).

In addition to be able to activate the classical pathway, CD40 is a strong inducer of the alternative NF-κB activating pathway. CD40 constitutive activation of B-cells increases IL-10 expression (8). As mentioned above, IL-10 gene expression was co-clusterized with the one of PD-L1 and PD-L2, being up-regulated in L.CD40 B-cell lymphoma (Figure 1A). IL-10 receptor signaling is mediated by the JAK1 and Tyk2 tyrosine kinase that leads to activation of STAT3 transcription factor, STAT3 being a major inducer of PD-L1 gene expression (9). CD40 stimulation can also augment BCR-induced B cell responses by activation of Bruton’s tyrosine kinase (BTK) (10), and the BCR is also able to induce the IL-10/STAT3 signaling with increased expression of PD-L1 in diffuse large B-cell lymphoma (9). We thus *ex vivo* treated purified L.CD40 lymphoma B-cells with the PHA-408 molecule, an inhibitor of IKK2/NF-κB activation, the JAK1/JAK2 tyrosine kinase inhibitor (TKI) ruxolitinib, and the BTK inhibitor ibrutinib. As shown in Figure 2, PD-L1 expression was strongly reduced 48h after treatment with either PHA-408 or ruxolitinib and moderately with ibrutinib, a strong indication that PD-L1 expression was under control of CD40 activation, either directly through NF-κB activation or indirectly through IL-10 induction of BCR sensitization and BTK activation.

**Figure 2.**
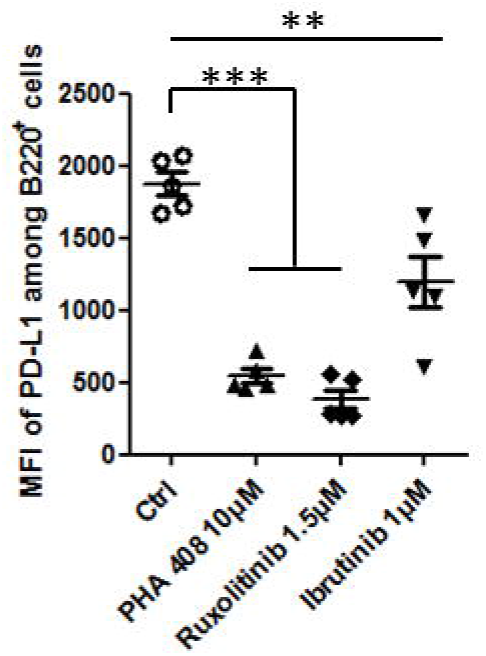
Flow cytometry Mean Fluorescence Intensity (MFI) of PD-L1 on B-cells after 48h *in vitro* inhibitor treatments (PHA 408, Ruxolitinib and Ibrutinib). Statistical significance was determined by unpaired t-test (***P<0.001; **P<0.01).

To address *in vivo* the role of PD-L1 in these B-cell lymphoma, L.CD40 mice were treated with an antibody blocking programmed cell death protein 1 (PD-1)/PD-1 1igand 1 (PD-L1) signaling for three weeks according to a methodology already described (11,12). We observed a reduction of spleen size and absolute number of splenocytes in anti-PD-L1 treated L.CD40 mice (Figure 3A), due to decreased B-cell numbers (Figure 3B). With PD-L1 blockade, we noticed a decrease in numbers of activated B-cells in L.CD40 mice (Figure 3C). This was associated with a reduction in the *in vivo* proliferation rate of spleen B-cells, as assessed by the decrease of *in vivo* BrdU incorporation over 18 hours (Figure 3D). Morphologically, spleen lymphocytes from anti-PD-L1 treated L.CD40 mice were smaller with a more condensed chromatin (Figure 3E). We then studied the impact of PD1/PD-L1 blockade in T-cell compartment. Expression of T-cell activation markers CD62L and CD44 was increased on both CD4^+^ and CD8^+^ T-cells (Figure 4A). In parallel, an increase in PD-1 expression was observed on the surface of CD4 T-cells. This increase in PD-1 expression was more heterogeneous on CD8 T-cells (Figure 4B).

**Figure 3.**
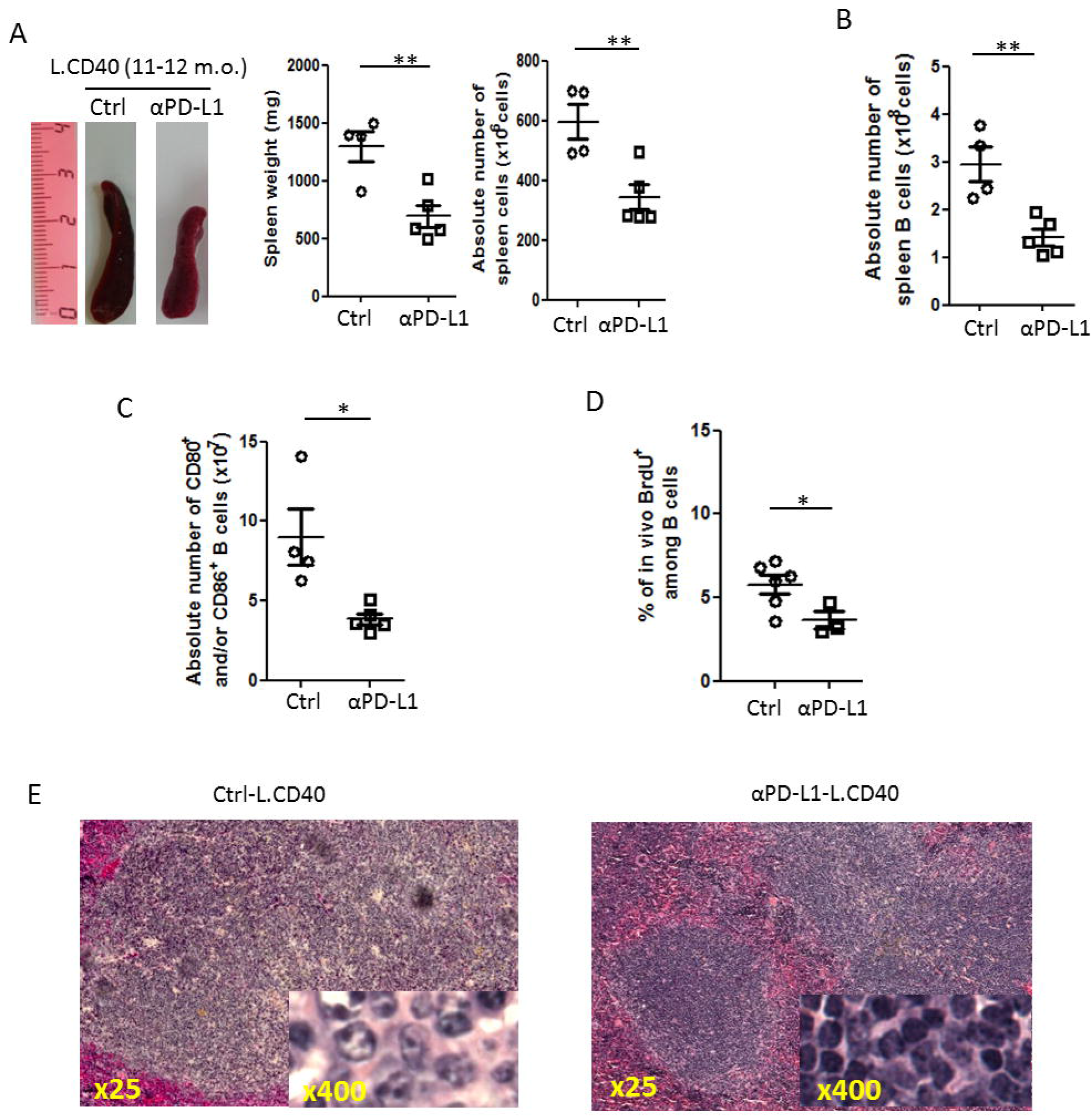
(A) Left panel, examples of whole spleens from Ctrl and αPD-L1 treated L.CD40 mice; middle panel, mean and standard deviation of spleen weight; right panel, absolute numbers of spleen cells. For the PD-L1 treatment, L.CD40 mice were injected every 4 days for three weeks with 200 μg anti-PD-L1 antibody in In VivoPure Dilution Buffer (clone 10F.9G2; Bio X cell; US). (B) Flow cytometry of spleen B220 B-cell absolute numbers in Ctrl and αPD-L1 treated L.CD40 mice. (C) Flow cytometry of absolute numbers of spleen B220 B-cells expressing CD80 and/or CD86 activation markers in L.CD40 after injection of isotope control (Ctrl) or anti-PD-L1 (αPD-L1) antibody. (D) Mean and standard deviation of flow cytometry percentages of BrdU positive B-cells after *in vivo* BrdU incorporation in Ctrl and αPD-L1 treated L.CD40 mice. Mice were injected intraperitoneally with 2 mg BrdU, 18 hours before isolating cells. (E) May Grunwald staining of spleen imprints of Ctrl-L.CD40 (left panel) and αPD-L1-L.CD40 (right panel) mice. Statistical significance was determined by unpaired t-test (**P<0.01; *P<0.05).

**Figure 4.**
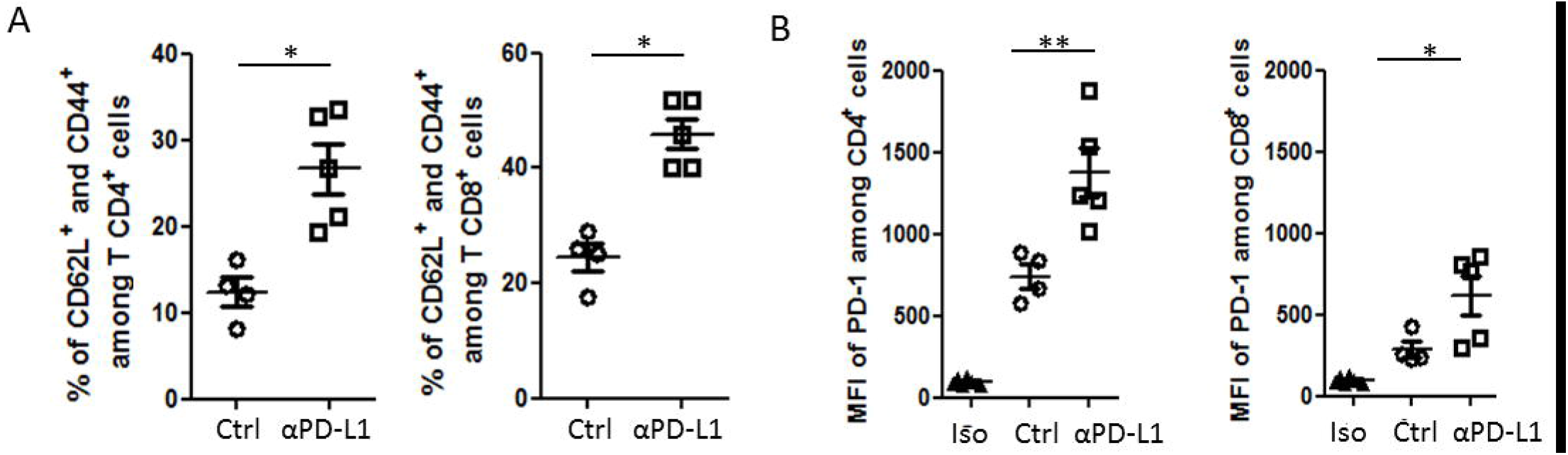
(A) Mean and standard deviation of flow cytometry percentages of activated CD4^+^ and CD8^+^ T cells expressing CD62L and CD44 activation markers in L.CD40 after injection of isotope control (Ctrl) or anti-PD-L1 (αPD-L1) antibody. (B) Flow cytometry Mean Fluorescence Intensity (MFI) of PD1 on CD4^+^ and CD8^+^ T cells in Ctrl and αPD-L1 treated L.CD40 mice. Statistical significance was determined by unpaired t-test (**P<0.01; *P<0.05).

## Discussion

Expression of PD-1 and PD-L1 in B-cell lymphomas and effect of immune therapies against the immune checkpoint axe has been recently reviewed (3). Expression of PD-L1 by tumor cells and effect of anti-PD-1 immune therapy has been clearly demonstrated in Hodgkin’s lymphoma, a tumor which is constantly associated with both NF-κB and STAT3 activation. In DLBCL, expression of PD-L1 is variable but participates to the gene immune escape signature of ABC-DLBCLs (3,7). ABC-DLBCL are not only associated with NF-κB activation but may exhibit a chronic active BCR (13) and are sensitive to BTK inhibitors (14). STAT3 activation mainly found in ABC-DLBCLs and is associated with poor survival (15).

Here, our results show that NF-κB activated B-cell lymphoma of the L.CD40 mouse model exhibited an immune escape gene signature involving expression of PD-L1 and PD-L2, which expression was co-clusterized with IL-10. Over-expression of PD-L1 expression not only involved NF-κB activation by CD40 but also BTK and JAK/STAT signaling, the latter probably being indirectly regulated via an autocrine loop with participation of IL-10 for example. Indeed, PD-L1 expression could be down-regulated after treated with NF-κB, JAK1/JAK2 and BTK inhibitors. Expression of PD-L1 was very likely to be associated to tumor immune escape as demonstrated for numerous solid cancers such as melanoma of lung cancers. Indeed, *in vivo* blockade of PD-L1 was able to rapidly repress expansion of these B-cell lymphomas, with concomitant decrease in both B-cell proliferation and B-cell expression of activation markers as well as an increase in T-cell activation. This clearly indicates that therapies against the PD-L1/PD-1 axis may work in lymphomas as long as the tumor cells express PD-L1. Our results also suggest that combination of immune therapy targeting the PD-1/PD-L1 axis and TKI specific for the JAK/STAT or the BCR/BTK pathway could be of interest, opening new perspective on the effect of these molecules on the reactivation of the immune system.

## Material and Methods

### Mouse models and *in vivo* and *ex vivo* treatments

L.CD40 mice have been already described (5). Animals were housed at 21–23°C with a 12-h light/dark cycle. All procedures were conducted under an approved protocol according to European guidelines for animal experimentation (French national authorization number: 87–022 and French ethics committee registration number “CREEAL”: 09–07-2012). For in vivo PD-L1 treatment, L.CD40 mice were injected intraperitoneally every 4 days for three weeks with 200 μg anti-PD-L1 antibody (clone 10F.9G2; Bio X cell; US). For ex vivo treatments, splenocytes were cultured for 48 hours in complete RPMI medium (Eurobio) supplemented with 10 % of FBS, 2mM of L-Glutamine, 1% of Na pyruvate, 100U/ml of penicillin and 100μg/ml of streptomycin (ThermoFisher Scientific) and with the following treatments: either 10 μM of PHA 408 or 1.5μM of Roxolitinib or 1μM of Ibrutinib.

### Flow cytometry

Spleen from mice were collected and immune cells were filtered through a sterile nylon membrane. Cell suspensions were stained at 4°C in FACS Buffer (PBS, 1% FBS, 2 mM EDTA) with the following fluorescent-labelled antibodies: anti B220-BV421 (clone RA3–6B2, 1/400), anti CD4-FITC (clone RM4-5, 1/2000), anti CD8a-APC (clone 53-6.7, 1/400), anti CD62L-BV421 (clone MEL-14, 1/200), anti CD44-PE (clone IM7, 1/200), anti-PD-L1-PE (clone 10F.9G2, 1/80), anti PD-1-PECy7 (clone 29F.1A12, 1/50), anti CD80-APC (clone16-10A1, 1/2500) and anti CD86-FITC (clone GL-1, 1/600). Stained cells were analyzed using a BD-Fortessa SORP flow cytometer (BD Bioscience; US). Results were analyzed using Kaluza Flow Cytometry software 1.2 (Beckman Coulter; France).

### Proliferation

For *in vivo* proliferation, mice were injected intraperitoneally with 2 mg BrdU (Sigma-Aldrich, US), 18 hours before isolating cells. Splenocytes were stained for B220 and phases of cell cycle were analyzed by measuring BrdU and Propidium Iodide (PI)-incorporation, using the FITC-BrdU Flow Kit (BD Pharmingen; US).

## Acknowledgments

We thank Dr J Cook Moreau, UMR CNRS 7276, Limoges, France, for English editing.

## Authorship Contributions

C.V.F. performed, analyzed experiments and contributed to the writing. L.R. helped to perform proliferation and flow cytometry experiments. U.Z.S. helped analyze the results and contributed to the writing of the manuscript. J.F. and N.F. designed and directed the study, contributed to the experiments, analyzed the results and wrote the manuscript.

