## Supplemental table 1 and 2 for "Pre-Clinical Blocking of PD-L1 molecule, which expression is down regulated by NF-κB, JAK1/JAK2 and BTK inhibitors, induces regression of activated B-cell lymphoma"

**Supplemental Table 1: Immune Escape Gene signature adapted to the Affymetrix HG-U133_Plus_2 array.** Probe annotations were retrieved with the AffyCompatible R package.

| **Affymetrix UniqId** | **NAME** | **SYMBOL** |
| --- | --- | --- |
| 206177_s_at | arginase, liver | ARG1 |
| 231662_at |  |  |
| 231663_s_at |  |  |
| 231665_at |  |  |
| 236226_at | B and T lymphocyte associated | BTLA |
| 216598_s_at | chemokine (C-C motif) ligand 2 | CCL2 |
| 207861_at | chemokine (C-C motif) ligand 22 | CCL22 |
| 203645_s_at | CD163 molecule | CD163 |
| 215049_x_at |  |  |
| 216233_at |  |  |
| 223834_at | CD274 molecule | CD274 |
| 227458_at |  |  |
| 1554519_at | CD80 molecule | CD80 |
| 1555689_at |  |  |
| 207176_s_at |  |  |
| 205685_at | CD86 molecule | CD86 |
| 205686_s_at |  |  |
| 210895_s_at |  |  |
| 204636_at | collagen, type XVII, alpha 1 | COL17A1 |
| 207082_at | colony stimulating factor 1 (macrophage) | CSF1 |
| 209716_at |  |  |
| 210557_x_at |  |  |
| 211839_s_at |  |  |
| 221331_x_at | cytotoxic T-lymphocyte-associated protein 4 | CTLA4 |
| 231794_at |  |  |
| 234362_s_at |  |  |
| 234895_at |  |  |
| 236341_at |  |  |
| 207691_x_at | ectonucleoside triphosphate diphosphohydrolase 1 | ENTPD1 |
| 209473_at |  |  |
| 209474_s_at |  |  |
| 228585_at |  |  |
| 221333_at | forkhead box P3 | FOXP3 |
| 221334_s_at |  |  |
| 224211_at |  |  |
| 221576_at | growth differentiation factor 15 | GDF15 |
| 221577_x_at |  |  |
| 229868_s_at |  |  |
| 1554285_at | hepatitis A virus cellular receptor 2 | HAVCR2 |
| 1555628_a_at | |  |
| 1555629_at |  |  |
| 235458_at |  |  |
| 209960_at | hepatocyte growth factor (hepapoietin A; scatter factor) | HGF |
| 209961_s_at |  |  |
| 210755_at |  |  |
| 210997_at |  |  |
| 210998_s_at |  |  |
| 210439_at | inducible T-cell co-stimulator | ICOS |
| 210029_at | indoleamine-pyrrole 2,3 dioxygenase | IDO1 |
| 1568638_a_at | indoleamine-pyrrole 2,3 dioxygenase-like 1 | IDO2 |
| 207433_at | interleukin 10 | IL10 |
| 207844_at | interleukin 13 | IL13 |
| 207538_at | interleukin 4 | IL4 |
| 207539_s_at |  |  |
| 205207_at | interleukin 6 (interferon, beta 2) | IL6 |
| 204863_s_at | interleukin 6 signal transducer (gp130, oncostatin M receptor) /// melanoma antigen family A, 4 | IL6ST |
| 204864_s_at |  |  |
| 211000_s_at |  |  |
| 212195_at |  |  |
| 212196_at |  |  |
| 234474_x_at |  |  |
| 234967_at |  |  |
| 1562031_at | Janus kinase 2 (a protein tyrosine kinase) | JAK2 |
| 205841_at |  |  |
| 205842_s_at |  |  |
| 208179_x_at | killer cell immunoglobulin-like receptor, two domains, long cytoplasmic tail, 3 | KIR2DL3 |
| 206486_at | lymphocyte-activation gene 3 | LAG3 |
| 208071_s_at | leukocyte-associated immunoglobulin-like receptor 1 | LAIR1 |
| 210644_s_at |  |  |
| 201105_at | lectin, galactoside-binding, soluble, 1 (galectin 1) | LGALS1 |
| 1557197_a_at | Lectin, galactoside-binding, soluble, 3 (galectin 3) | LGALS3 |
| 208949_s_at |  |  |
| 200796_s_at | myeloid cell leukemia sequence 1 (BCL2-related) | MCL1 |
| 200797_s_at |  |  |
| 200798_x_at |  |  |
| 214056_at |  |  |
| 214057_at |  |  |
| 227175_at |  |  |
| 204438_at | mannose receptor, C type 1 /// mannose receptor, C type 1-like 1 | MRC1 |
| 208422_at | macrophage scavenger receptor 1 | MSR1 |
| 208423_s_at |  |  |
| 211887_x_at |  |  |
| 214770_at |  |  |
| 202431_s_at | v-myc myelocytomatosis viral oncogene homolog (avian) | MYC |
| 1553994_at | 5'-nucleotidase, ecto (CD73) | NT5E |
| 1553995_a_at | |  |
| 203939_at |  |  |
| 227486_at |  |  |
| 207634_at | programmed cell death 1 | PDCD1 |
| 220049_s_at | programmed cell death 1 ligand 2 | PDCD1LG2 |
| 224399_at |  |  |
| 209193_at | pim-1 oncogene /// pim-1 oncogene | PIM1 |
| 205127_at | prostaglandin-endoperoxide synthase 1 (prostaglandin G/H synthase and cyclooxygenase) | PTGS1 |
| 205128_x_at |  |  |
| 215813_s_at |  |  |
| 238669_at |  |  |
| 1554997_a_at | prostaglandin-endoperoxide synthase 2 (prostaglandin G/H synthase and cyclooxygenase) | PTGS2 |
| 204748_at |  |  |
| 1556582_at | CDNA FLJ25946 fis, clone JTH14258 | PVR |
| 212662_at |  |  |
| 214443_at |  |  |
| 214444_s_at |  |  |
| 216283_s_at |  |  |
| 239918_at |  |  |
| 32699_s_at |  |  |
| 206359_at | suppressor of cytokine signaling 3 | SOCS3 |
| 206360_s_at |  |  |
| 214105_at |  |  |
| 227697_at |  |  |
| 208991_at | signal transducer and activator of transcription 3 (acute-phase response factor) | STAT3 |
| 208992_s_at |  |  |
| 225289_at |  |  |
| 243213_at |  |  |
| 203084_at | transforming growth factor, beta 1 (Camurati-Engelmann disease) | TGFB1 |
| 203085_s_at |  |  |
| 240070_at | V-set and immunoglobulin domain containing 9 | TIGIT |
| 201666_at | TIMP metallopeptidase inhibitor 1 | TIMP1 |
| 209354_at | tumor necrosis factor receptor superfamily, member 14 (herpesvirus entry mediator) | TNFRSF14 |
| 210512_s_at | vascular endothelial growth factor | VEGFA |
| 210513_s_at |  |  |
| 211527_x_at |  |  |
| 212171_x_at |  |  |
| 219768_at | V-set domain containing T cell activation inhibitor 1 | VTCN1 |

**Supplemental Table 2: Immune Escape Gene signature adapted to the HT MG-430 PM Array.** Probe annotations were retrieved with the AffyCompatible R package.

| **Affymetrix UniqId** | **NAME** | **SYMBOL** |
| --- | --- | --- |
| 1419549_PM_at | arginase, liver | ARG1 |
| 1455656_PM_at | B and T lymphocyte associated | BTLA |
| 1420380_PM_at | chemokine (C-C motif) ligand 2 | CCL2 |
| 1417925_PM_at | chemokine (C-C motif) ligand 22 | CCL22 |
| 1419144_PM_at | CD163 antigen | CD163 |
| 1419714_PM_at | CD274 antigen | CD274 |
| 1427717_PM_at | CD80 antigen | CD80 |
| 1432826_PM_a_at | |  |
| 1451950_PM_a_at | |  |
| 1454372_PM_at | |  |
| 1420404_PM_at | CD86 antigen | CD86 |
| 1449858_PM_at | |  |
| 1418799_PM_a_at | collagen, type XVII, alpha 1 | COL17A1 |
| 1425154_PM_a_at | colony stimulating factor 1 (macrophage) | CSF1 |
| 1425155_PM_x_at | |  |
| 1448914_PM_a_at | |  |
| 1460220_PM_a_at | |  |
| 1419334_PM_at | cytotoxic T-lymphocyte-associated protein 4 | CTLA4 |
| 1423326_PM_at | ectonucleoside triphosphate diphosphohydrolase 1 | ENTPD1 |
| 1450939_PM_at | |  |
| 1420765_PM_a_at | forkhead box P3 | FOXP3 |
| 1418949_PM_at | growth differentiation factor 15 | GDF15 |
| 1451584_PM_at | hepatitis A virus cellular receptor 2 | HAVCR2 |
| 1425379_PM_at | hepatocyte growth factor | HGF |
| 1442884_PM_at | |  |
| 1451866_PM_a_at | |  |
| 1458943_PM_at | --- | HGF |
| 1421930_PM_at | inducible T cell co-stimulator | ICOS |
| 1421931_PM_at | |  |
| 1436598_PM_at | |  |
| 1420437_PM_at | indoleamine 2,3-dioxygenase 1 | IDO1 |
| 1425778_PM_at | indoleamine 2,3-dioxygenase 2 | IDO2 |
| 1450330_PM_at | interleukin 10 | IL10 |
| 1420802_PM_at | interleukin 13 | IL13 |
| 1449864_PM_at | interleukin 4 | IL4 |
| 1450297_PM_at | interleukin 6 | IL6 |
| 1421239_PM_at | interleukin 6 signal transducer | IL6ST |
| 1437303_PM_at | |  |
| 1452843_PM_at | |  |
| 1460295_PM_s_at | |  |
| 1421065_PM_at | Janus kinase 2 | JAK2 |
| 1421066_PM_at | |  |
| 1449911_PM_at | lymphocyte-activation gene 3 | LAG3 |
| 1430447_PM_a_at | leukocyte-associated Ig-like receptor 1 | LAIR1 |
| 1439067_PM_at | |  |
| 1444040_PM_at | |  |
| 1419573_PM_a_at | lectin, galactose binding, soluble 1 | LGALS1 |
| 1455439_PM_a_at | |  |
| 1426808_PM_at | lectin, galactose binding, soluble 3 | LGALS3 |
| 1416880_PM_at | myeloid cell leukemia sequence 1 | MCL1 |
| 1416881_PM_at | |  |
| 1437527_PM_x_at | |  |
| 1448503_PM_at | |  |
| 1456243_PM_x_at | |  |
| 1456381_PM_x_at | |  |
| 1450430_PM_at | mannose receptor, C type 1 | MRC1 |
| 1422062_PM_at | macrophage scavenger receptor 1 | MSR1 |
| 1425434_PM_a_at | |  |
| 1425435_PM_at | |  |
| 1448061_PM_at | |  |
| 1424942_PM_a_at | myelocytomatosis oncogene | MYC |
| 1422974_PM_at | 5' nucleotidase, ecto | NT5E |
| 1428547_PM_at | |  |
| 1459023_PM_at | |  |
| 1449835_PM_at | programmed cell death 1 | PDCD1 |
| 1450290_PM_at | programmed cell death 1 ligand 2 | PDCD1LG2 |
| 1423006_PM_at | proviral integration site 1 | PIM1 |
| 1435458_PM_at | |  |
| 1423414_PM_at | prostaglandin-endoperoxide synthase 1 | PTGS1 |
| 1436448_PM_a_at | |  |
| 1417262_PM_at | prostaglandin-endoperoxide synthase 2 | PTGS2 |
| 1417263_PM_at | |  |
| 1423903_PM_at | poliovirus receptor | PVR |
| 1423904_PM_a_at | |  |
| 1423905_PM_at | |  |
| 1429848_PM_at | |  |
| 1450295_PM_s_at | |  |
| 1451160_PM_s_at | |  |
| 1416576_PM_at | suppressor of cytokine signaling 3 | SOCS3 |
| 1455899_PM_x_at | |  |
| 1456212_PM_x_at | |  |
| 1424272_PM_at | signal transducer and activator of transcription 3 | STAT3 |
| 1426587_PM_a_at | |  |
| 1460700_PM_at | |  |
| 1420653_PM_at | transforming growth factor, beta 1 | TGFB1 |
| 1460227_PM_at | tissue inhibitor of metalloproteinase 1 | TIMP1 |
| 1452425_PM_at | tumor necrosis factor receptor superfamily, member 14 (herpesvirus entry mediator) | TNFRSF14 |
| 1420909_PM_at | vascular endothelial growth factor A | VEGFA |
| 1451959_PM_a_at | |  |
| 1458070_PM_at | V-set domain containing T cell activation inhibitor 1 | VTCN1 |
