## Supplemental table 3 for "Pre-Clinical Blocking of PD-L1 molecule, which expression is down regulated by NF-κB, JAK1/JAK2 and BTK inhibitors, induces regression of activated B-cell lymphoma"

| Gene.Symbol | SYMBOL.OK | S1 | S2 | S3 | S4 | S5 | S6 | S7 | S8 | S9 | S10 | S11 |
| --- | --- | --- | --- | --- | --- | --- | --- | --- | --- | --- | --- | --- |
|  |  | L.CD40 | L.CD40 | WT | L.CD40 | WT | L.CD40 | WT | WT | L.CD40 | L.CD40 | WT |
| Socs3 | SOCS3 | 6,005909 | 7,00368 | 6,21625 | 5,679432 | 6,639841 | 5,576802 | 5,660009 | 5,79407 | 5,882332 | 6,715188 | 6,119701 |
| Mcl1 | MCL1 | 10,508 | 10,49781 | 10,46064 | 10,54331 | 10,36326 | 10,57337 | 10,29679 | 10,31036 | 10,55534 | 10,49081 | 10,49446 |
| Mcl1 | MCL1 | 9,46207 | 9,56691 | 9,227069 | 9,294104 | 8,939931 | 9,805619 | 9,400434 | 9,188602 | 9,321149 | 9,469131 | 9,267929 |
| Ptgs2 | PTGS2 | 3,3812 | 7,468712 | 4,386524 | 3,154966 | 4,305453 | 3,048162 | 2,919847 | 2,52821 | 2,951494 | 7,825598 | 4,700265 |
| Ptgs2 | PTGS2 | 2,589067 | 3,804557 | 2,862831 | 2,796341 | 2,867722 | 2,514885 | 2,83243 | 2,914149 | 2,643327 | 3,933054 | 2,986423 |
| Ccl22 | CCL22 | 7,863233 | 7,901059 | 8,492901 | 7,544222 | 8,71841 | 7,304679 | 8,185423 | 8,24209 | 7,745826 | 7,821568 | 8,382617 |
| Col17a1 | COL17A1 | 3,226818 | 3,459077 | 3,398368 | 3,431578 | 3,692775 | 3,185287 | 3,239369 | 3,343516 | 3,368347 | 3,404559 | 3,307892 |
| Gdf15 | GDF15 | 4,252857 | 4,608806 | 4,488442 | 4,438666 | 4,235334 | 4,16476 | 4,157464 | 3,992729 | 4,180928 | 4,674397 | 4,457056 |
| Cd163 | CD163 | 3,479063 | 3,128074 | 3,649386 | 3,391378 | 3,834991 | 3,688342 | 3,683553 | 4,213252 | 3,231764 | 3,261976 | 3,901889 |
| Ctla4 | CTLA4 | 6,566883 | 7,103453 | 7,150558 | 6,762147 | 7,039824 | 6,20697 | 7,261214 | 7,409065 | 6,667328 | 7,105734 | 7,088212 |
| Arg1 | ARG1 | 2,479284 | 2,499345 | 2,46396 | 2,496854 | 2,401689 | 2,492336 | 2,336485 | 2,575417 | 2,496122 | 2,507623 | 2,421262 |
| Lgals1 | LGALS1 | 9,665716 | 9,878941 | 9,856366 | 9,484958 | 9,551316 | 8,839795 | 9,097919 | 9,179095 | 9,699948 | 9,704531 | 9,987165 |
| Cd274 | CD274 | 7,725136 | 7,578328 | 7,18901 | 7,664454 | 7,275939 | 7,74213 | 7,633124 | 7,421613 | 7,610918 | 7,541961 | 7,194985 |
| Ccl2 | CCL2 | 6,245352 | 9,243315 | 7,830642 | 5,338832 | 6,320584 | 4,954648 | 5,504246 | 5,187937 | 6,347934 | 9,048187 | 7,750871 |
| Cd86 | CD86 | 8,313865 | 8,441431 | 8,30098 | 8,123225 | 8,249434 | 8,25216 | 8,343554 | 8,296757 | 8,281857 | 8,311126 | 8,273252 |
| Ido1 | IDO1 | 2,931803 | 3,061256 | 3,102196 | 2,890253 | 3,062946 | 3,176277 | 2,918091 | 2,925395 | 2,801868 | 3,029672 | 2,869917 |
| Tgfb1 | TGFB1 | 5,707906 | 5,070099 | 4,734338 | 4,696856 | 5,320621 | 4,489656 | 4,552443 | 4,780471 | 4,810358 | 5,128934 | 4,900513 |
| Foxp3 | FOXP3 | 4,482437 | 4,469159 | 4,399957 | 4,377188 | 4,709731 | 3,991521 | 4,189839 | 3,984134 | 4,284181 | 4,369176 | 4,438973 |
| Il13 | IL13 | 3,55499 | 3,526098 | 3,617186 | 3,587657 | 3,517996 | 3,261895 | 3,421223 | 3,385197 | 3,591142 | 3,635904 | 3,476151 |
| Vegfa | VEGFA | 6,284728 | 6,195731 | 6,472678 | 6,018229 | 6,083447 | 5,96968 | 5,927129 | 5,750267 | 6,297006 | 6,421124 | 6,447956 |
| Jak2 | JAK2 | 7,403181 | 7,435638 | 7,355487 | 7,60665 | 7,194346 | 7,493603 | 7,377897 | 7,301309 | 7,450103 | 7,391317 | 7,338479 |
| Jak2 | JAK2 | 8,827711 | 8,88147 | 8,822062 | 9,0914 | 8,861868 | 8,998153 | 8,851313 | 8,96769 | 8,839447 | 8,919144 | 8,930855 |
| Il6st | IL6ST | 4,475723 | 4,543486 | 4,439378 | 4,395066 | 4,379219 | 4,594856 | 4,469586 | 4,49241 | 4,106584 | 4,511612 | 4,273371 |
| Icos | ICOS | 7,002573 | 7,058133 | 7,316708 | 6,770706 | 7,868952 | 7,248888 | 7,496714 | 7,496599 | 7,093966 | 7,2632 | 7,099194 |
| Icos | ICOS | 5,547511 | 5,674193 | 5,689719 | 5,269573 | 5,749516 | 5,579022 | 6,040211 | 5,846566 | 5,507093 | 5,634839 | 5,28941 |
| Msr1 | MSR1 | 4,910955 | 5,098832 | 4,787189 | 5,048219 | 4,490539 | 5,016678 | 5,597762 | 5,239937 | 4,934818 | 5,171609 | 4,710237 |
| Nt5e | NT5E | 4,504411 | 4,67392 | 4,610262 | 4,58913 | 4,441724 | 4,653879 | 4,558133 | 4,854476 | 4,547909 | 4,492916 | 4,518311 |
| Pim1 | PIM1 | 4,817733 | 5,058193 | 4,747154 | 5,391355 | 4,822171 | 5,011425 | 4,886978 | 4,975605 | 4,812928 | 4,928475 | 4,770113 |
| Entpd1 | ENTPD1 | 5,723202 | 5,888308 | 5,446023 | 5,775379 | 5,133922 | 6,164463 | 5,234172 | 5,554892 | 5,748389 | 5,786559 | 5,674716 |
| Ptgs1 | PTGS1 | 5,156952 | 4,738912 | 5,207957 | 5,081603 | 5,017428 | 4,875799 | 4,918845 | 5,125225 | 5,21488 | 4,987503 | 5,07742 |
| Pvr | PVR | 5,743432 | 5,794309 | 5,889822 | 5,623197 | 5,742644 | 5,506263 | 5,454568 | 5,690054 | 5,698403 | 5,674073 | 5,90057 |
| Pvr | PVR | 6,24994 | 6,545319 | 6,49126 | 6,357008 | 6,40266 | 5,812297 | 6,221028 | 6,061428 | 6,046543 | 6,23331 | 6,538468 |
| Pvr | PVR | 5,103445 | 5,325464 | 5,429975 | 4,828321 | 4,737517 | 4,63338 | 4,936173 | 4,735725 | 4,862772 | 5,306854 | 5,165046 |
| Stat3 | STAT3 | 4,481663 | 4,660243 | 4,598126 | 4,466928 | 4,564013 | 4,939942 | 4,614423 | 4,753909 | 4,638445 | 4,652137 | 4,550607 |
| Myc | MYC | 8,198873 | 8,881454 | 8,134664 | 7,930717 | 8,591174 | 7,182569 | 7,57096 | 7,690895 | 8,229244 | 8,473869 | 8,135545 |
| Csf1 | CSF1 | 4,342192 | 4,796363 | 4,368155 | 4,593549 | 4,119582 | 4,286386 | 4,059582 | 4,16486 | 4,374975 | 4,340852 | 4,556793 |
| Csf1 | CSF1 | 4,470396 | 4,116882 | 4,337021 | 4,60033 | 4,651237 | 4,068852 | 3,916633 | 4,323425 | 4,223038 | 4,279936 | 4,542304 |
| Hgf | HGF | 3,602359 | 3,486483 | 3,704237 | 3,72647 | 3,717575 | 3,686913 | 3,779844 | 3,675767 | 3,481374 | 3,691589 | 3,512272 |
| Msr1 | MSR1 | 4,035295 | 3,902386 | 4,014551 | 4,366845 | 3,773847 | 3,823112 | 4,251291 | 3,979804 | 4,08977 | 4,043066 | 4,002342 |
| Msr1 | MSR1 | 2,603135 | 2,644112 | 2,745285 | 2,750055 | 2,92363 | 2,944405 | 2,706864 | 2,843593 | 2,68395 | 2,609145 | 2,538946 |
| Ido2 | IDO2 | 3,284227 | 3,184 | 3,545616 | 3,211224 | 3,099673 | 3,15495 | 3,396286 | 3,085055 | 3,229701 | 3,235199 | 3,273399 |
| Stat3 | STAT3 | 8,346646 | 8,519179 | 8,539443 | 8,00066 | 8,424967 | 8,088336 | 8,343226 | 8,367154 | 8,398946 | 8,358907 | 8,253231 |
| Lgals3 | LGALS3 | 9,087753 | 9,175596 | 9,166606 | 9,004145 | 8,985965 | 9,112946 | 9,392373 | 9,371975 | 9,104593 | 9,160115 | 9,076097 |
| Cd80 | CD80 | 4,280097 | 4,639936 | 4,413362 | 4,476752 | 4,391054 | 4,632593 | 4,268074 | 4,126484 | 4,223114 | 4,818595 | 4,454071 |
| Nt5e | NT5E | 5,558236 | 6,118557 | 5,965364 | 5,986161 | 5,706074 | 6,08232 | 6,307717 | 6,391449 | 5,669964 | 6,131616 | 6,001642 |
| Pvr | PVR | 2,861013 | 3,091856 | 3,033509 | 3,107512 | 3,088794 | 3,011459 | 3,042726 | 3,054276 | 3,174016 | 2,948874 | 2,745538 |
| Lair1 | LAIR1 | 5,550704 | 5,538179 | 5,706176 | 5,907406 | 5,597755 | 5,795677 | 5,934569 | 5,732846 | 5,661344 | 5,462294 | 5,835833 |
| Cd80 | CD80 | 4,914718 | 4,992198 | 4,734995 | 4,622925 | 4,138494 | 4,676195 | 4,466939 | 4,349532 | 4,821135 | 4,990409 | 4,768587 |
| Pim1 | PIM1 | 7,581176 | 7,697744 | 7,634966 | 8,617551 | 7,604256 | 7,589149 | 7,661356 | 7,343645 | 7,654653 | 7,674097 | 7,537831 |
| Ptgs1 | PTGS1 | 3,897074 | 3,803078 | 3,843559 | 3,84919 | 3,766288 | 3,577651 | 3,909493 | 4,258012 | 3,810627 | 3,702881 | 3,952111 |
| Icos | ICOS | 8,703886 | 8,944343 | 9,009409 | 8,712566 | 8,948844 | 8,438395 | 9,067164 | 8,858515 | 8,552727 | 8,868976 | 8,995528 |
| Il6st | IL6ST | 7,55444 | 7,520274 | 7,961209 | 7,416751 | 8,85911 | 7,365457 | 7,161427 | 7,101431 | 7,652895 | 7,620235 | 7,645761 |
| Mcl1 | MCL1 | 11,35513 | 11,39623 | 11,34581 | 11,36104 | 11,28711 | 11,37505 | 11,26067 | 11,27367 | 11,39498 | 11,39703 | 11,42619 |
| Lair1 | LAIR1 | 4,398982 | 4,47199 | 4,804315 | 4,419465 | 4,82786 | 4,605865 | 4,497637 | 4,508711 | 4,611871 | 4,324157 | 4,562533 |
| Hgf | HGF | 3,237302 | 3,198099 | 3,317764 | 3,307162 | 3,173371 | 3,023923 | 3,342925 | 3,192173 | 3,309876 | 3,247235 | 3,122883 |
| Lair1 | LAIR1 | 6,072849 | 5,85053 | 6,166424 | 6,145582 | 6,44243 | 5,939441 | 6,307034 | 6,107123 | 5,958361 | 5,96724 | 6,251432 |
| Msr1 | MSR1 | 6,51703 | 6,407607 | 6,281767 | 6,695505 | 6,175049 | 6,638416 | 7,400377 | 7,22543 | 6,421213 | 6,551739 | 6,385352 |
| Mcl1 | MCL1 | 11,36944 | 11,35611 | 11,19926 | 11,30367 | 11,0205 | 11,40442 | 11,20493 | 11,14765 | 11,23099 | 11,32218 | 11,19259 |
| Csf1 | CSF1 | 5,49036 | 5,369506 | 5,79743 | 5,462217 | 5,809564 | 5,928751 | 5,303217 | 5,460175 | 5,393423 | 5,557876 | 5,648861 |
| Pdcd1 | PDCD1 | 5,072269 | 5,561886 | 5,072433 | 5,086697 | 5,238043 | 4,667822 | 4,377559 | 4,7219 | 5,1227 | 5,453894 | 5,157772 |
| Cd86 | CD86 | 7,772103 | 8,064103 | 7,504749 | 7,765427 | 7,194769 | 8,087268 | 7,64436 | 7,477357 | 7,838511 | 8,026646 | 7,646186 |
| Il4 | IL4 | 3,16053 | 3,212999 | 3,047667 | 3,016475 | 2,939034 | 3,22308 | 3,136806 | 3,169046 | 3,144932 | 3,026935 | 2,977153 |
| Lag3 | LAG3 | 4,654988 | 5,200348 | 4,706776 | 4,595956 | 4,964516 | 4,574366 | 5,097552 | 4,964905 | 4,766283 | 4,593592 | 4,860339 |
| Pdcd1lg2 | PDCD1LG2 | 5,779895 | 5,912698 | 5,365527 | 6,460982 | 5,190663 | 6,136878 | 5,25691 | 4,982503 | 5,84876 | 5,817204 | 5,375277 |
| Pvr | PVR | 3,169854 | 3,301679 | 2,934589 | 2,85431 | 2,792347 | 3,134273 | 3,341067 | 3,130305 | 3,189097 | 3,026806 | 3,025108 |
| Il6 | IL6 | 2,738676 | 5,407804 | 2,837938 | 2,751302 | 3,005056 | 2,830559 | 2,923219 | 2,667493 | 2,865811 | 5,275456 | 2,794873 |
| Il10 | IL10 | 3,649813 | 3,633561 | 3,647704 | 3,670089 | 3,449393 | 3,653646 | 3,400966 | 3,665276 | 3,612764 | 3,7061 | 3,538469 |
| Mrc1 | MRC1 | 7,311781 | 7,072989 | 7,447702 | 7,761845 | 7,164617 | 7,10478 | 8,256294 | 8,389089 | 7,483661 | 7,054807 | 7,678877 |
| Entpd1 | ENTPD1 | 4,352449 | 4,576404 | 4,141538 | 4,310465 | 3,901995 | 4,321336 | 3,735402 | 4,106806 | 4,449742 | 4,308924 | 3,99922 |
| Pvr | PVR | 6,119404 | 6,416634 | 6,321443 | 6,040103 | 6,269042 | 5,976427 | 6,3234 | 6,359293 | 6,204997 | 6,208919 | 6,301095 |
| Havcr2 | HAVCR2 | 5,563991 | 5,468953 | 5,370176 | 5,382811 | 5,251351 | 5,206402 | 5,012296 | 5,147601 | 5,371295 | 5,382558 | 5,437147 |
| Hgf | HGF | 3,142769 | 3,260716 | 3,125634 | 3,185446 | 3,249965 | 2,881745 | 3,205103 | 3,205068 | 2,945383 | 3,046134 | 3,14615 |
| Cd80 | CD80 | 5,585298 | 5,735782 | 5,346452 | 5,301634 | 4,809873 | 5,779509 | 5,219222 | 5,102713 | 5,561255 | 5,790864 | 5,340969 |
| Vegfa | VEGFA | 4,039178 | 4,34825 | 4,005418 | 4,084904 | 3,94544 | 3,898549 | 3,947271 | 3,673937 | 4,075689 | 4,019248 | 3,922158 |
| Tnfrsf14 | TNFRSF14 | 6,665705 | 6,766764 | 6,66594 | 6,540808 | 6,736009 | 6,19431 | 4,099058 | 4,291727 | 6,569295 | 6,622931 | 6,576703 |
| Il6st | IL6ST | 8,936183 | 8,788777 | 9,107941 | 8,949299 | 9,066668 | 8,686652 | 9,366437 | 9,536213 | 8,91623 | 8,703604 | 9,066284 |
| Cd80 | CD80 | 3,368085 | 3,26341 | 3,38998 | 3,275187 | 3,226056 | 3,285347 | 3,18608 | 3,189101 | 3,358795 | 3,365033 | 3,275272 |
| Lgals1 | LGALS1 | 10,49598 | 10,53794 | 10,7245 | 10,40051 | 10,60613 | 10,0158 | 10,40274 | 10,43064 | 10,60697 | 10,58409 | 10,81073 |
| Btla | BTLA | 10,4549 | 10,52395 | 10,49819 | 10,63813 | 10,19578 | 11,01282 | 10,89751 | 11,28688 | 10,48722 | 10,54732 | 10,57087 |
| Socs3 | SOCS3 | 9,097577 | 10,24814 | 9,277944 | 8,858137 | 9,072134 | 8,391457 | 8,431899 | 8,312517 | 9,197945 | 9,9884 | 9,267189 |
| Socs3 | SOCS3 | 7,997905 | 8,886133 | 8,174846 | 7,832374 | 8,255231 | 6,986619 | 7,232568 | 7,129918 | 8,027341 | 8,742461 | 8,158438 |
| Mcl1 | MCL1 | 10,96253 | 10,9618 | 10,95972 | 10,95312 | 10,8904 | 11,03824 | 10,89222 | 10,8808 | 10,99656 | 10,96265 | 10,99404 |
| Mcl1 | MCL1 | 10,73028 | 10,78034 | 10,65825 | 10,71873 | 10,5781 | 10,33327 | 10,34231 | 10,30375 | 10,80496 | 10,72057 | 10,66581 |
| Vtcn1 | VTCN1 | 2,595066 | 2,697918 | 2,721091 | 2,609436 | 2,587138 | 2,642861 | 2,56887 | 2,480808 | 2,543681 | 2,676431 | 2,541723 |
| --- | HGF | 2,876143 | 2,734534 | 2,885118 | 2,801489 | 2,768417 | 2,810553 | 2,71796 | 2,761521 | 2,823497 | 2,915735 | 2,999007 |
| Nt5e | NT5E | 3,240899 | 3,333322 | 3,404814 | 3,373247 | 3,265854 | 3,400437 | 3,374316 | 3,084415 | 3,144768 | 3,247399 | 3,287018 |
| Csf1 | CSF1 | 7,045674 | 6,967881 | 7,128151 | 7,048606 | 7,140875 | 7,311327 | 6,962286 | 7,331048 | 6,957867 | 6,741194 | 6,921062 |
| Timp1 | TIMP1 | 4,212571 | 4,624496 | 4,528072 | 4,702196 | 4,743933 | 4,558552 | 4,730231 | 4,384658 | 4,560932 | 4,640856 | 4,476967 |
| Il6st | IL6ST | 6,050394 | 5,798968 | 5,974341 | 5,747455 | 5,148949 | 5,661009 | 6,615504 | 6,552331 | 5,916802 | 5,796712 | 6,071505 |
| Stat3 | STAT3 | 8,538157 | 8,675438 | 8,674687 | 8,422579 | 8,351134 | 8,194937 | 8,25638 | 8,483295 | 8,436516 | 8,49811 | 8,573049 |
